# Exenatide reverts the high-fat-diet-induced impairment of BDNF signaling and inflammatory response in an animal model of Alzheimer’s disease

**DOI:** 10.1101/487629

**Authors:** Manuela Bomba, Alberto Granzotto, Vanessa Castelli, Rossano Lattanzio, Annamaria Cimini, Stefano L. Sensi

**Affiliations:** Center of Excellence on Aging and Translational Medicine ‑ CeSI-MeT, University G. d’Annunzio of Chieti-Pescara, Italy; Department of Neuroscience, Imaging, and Clinical Sciences, University G. d’Annunzio of Chieti-Pescara, Italy; Department of Life, Health and Environmental Sciences, University of L’Aquila, Italy; Department of Medical, Oral, and Biotechnological Sciences, University G. d’Annunzio of Chieti-Pescara, Italy; Sbarro Institute for Cancer Research and Molecular Medicine and Center for Biotechnology, Temple University, Philadelphia, USA; National Institute for Nuclear Physics (INFN), Gran Sasso National Laboratory (LNGS), Assergi, Italy; Departments of Neurology and Pharmacology, Institute for Mind Impairments and Neurological Disorders - iMIND, University of California - Irvine, Irvine, USA

**Keywords:** exendin-4, insulin, diabetes, T2DM, insulin resistance, BDNF, neurotrophic factors, dementia, aging, synaptic plasticity, memory

## Abstract

**Background:** preclinical, clinical, and epidemiological evidence support the notion that Alzheimer’s disease(AD) is a multifactorial condition in which, along with β-amyloid (Aβ) and tau-related pathology, the synergistic activity of genetic, environmental, vascular, metabolic, and inflammatory factors promote the onset and progression of the disease. Epidemiological evidence indicate that glucose intolerance, deficits in insulin secretion or type 2 diabetes mellitus (T2DM) participate in increasing the risk of developing cognitive impairment or dementia. A pivotal role in the process is played by insulin as the hormone critically regulates brain functioning. GLP-1, the glucagon-like peptide 1, facilitates insulin signaling, regulates glucose homeostasis, and modulates synaptic plasticity. Exenatide is a GLP-1R agonist, characterized by an extended half-life, employed in T2DM. However, exenatide has also been shown to affect the signaling of the brain-derived neurotrophic factor (BDNF), synaptic plasticity, and cognitive performances in animal models of brain aging and neurodegeneration.

**Methods:** in this study, we tested whether exenatide exerts neuroprotection in a preclinical AD model set to mimic the clinical complexity of the human disease. To that aim, we investigated the effects of 3-month exenatide treatment in 3xTg-AD mice challenged for six months with a high-fat diet (HFD). Endpoints of the study were variations in systemic metabolism, insulin and neurotrophic signaling, neuroinflammation, levels of Aβ and tau pathology as well as changes in cognitive performances.

**Findings and interpretation:** results of the study indicate that exenatide reverts the adverse changes of BDNF signaling and the neuroinflammation status of 3xTg-AD mice undergoing HFD.

## Introduction

Alzheimer’s disease (AD) is a neurodegenerative condition associated with the presence of cognitive and behavioral deficits and characterized by the accumulation of neuritic plaques composed of β–amyloid (Aβ) peptides, the appearance of neurofibrillary tangles (NFT) made of hyperphosphorylated tau (p-tau) proteins, and reactive gliosis ^1^. Preclinical, clinical, and epidemiological data support the notion that AD is a multifactorial condition in which, along with Aβ and tau-related pathology, the convergence of genetic, environmental, vascular, metabolic, and inflammatory factors promotes the onset and development of the disease ^2–7^. In that regard, metabolic disorders are the objects of growing therapeutic attention ^8–11^. Epidemiological studies indicate that glucose intolerance, deficits in insulin secretion or type 2 diabetes (T2DM) increase the risk of developing cognitive impairment or dementia ^11^.

In the Central Nervous System (CNS), insulin regulates many critical biological events occurring in neurons, glia, microglia, and the neurovascular unit. In that compartments, insulin participates in the control of energy homeostasis, the modulation of protein and lipid synthesis, apoptosis, autophagy, the permeability of the blood-brain barrier, neurotransmitter balance, cytoskeletal remodeling, synaptic plasticity and neurogenesis ^11–13^. Insulin resistance (IR), a condition observed in T2DM patients, has also been described in the brain of AD patients. In the brain, IR can occur even in the absence of obesity, peripheral IR or overt signs of T2DM ^14^. In the CNS, IR leads to impaired structural and functional plasticity, ultimately contributing to the development of neuronal and brain dysfunctions ^11,15,16^.

GLP-1, the glucagon-like peptide 1, is a hormone that facilitates insulin signaling and regulates glucose homeostasis ^17,18^. However, GLP-1 receptors (GLP-1Rs) are also expressed in the brain ^17^ where they contribute to the modulation of neuronal excitability, synaptic plasticity, and cognition ^19–21^. Exenatide is a GLP-1R agonist, characterized by an extended half-life, and employed in T2DM. The molecule also promotes beneficial effects in the CNS ^3,22,23^. In preclinical models of brain aging and neurodegeneration, exenatide has been shown to positively affect the signaling of the brain-derived neurotrophic factor (BDNF) and to modulate synaptic plasticity and cognitive performances ^24–27^. In clinical settings, exenatide has been successfully employed to alleviate the motor symptoms of Parkinson’s disease (PD) patients ^28,29^.

In this study, we tested whether exenatide promotes neuroprotection in a preclinical AD model that was set to mimic the clinical complexity of the human disease. To that aim, we evaluated the effects of a 3-month treatment in 3xTg-AD mice that were challenged for six months with a high-fat diet (HFD). Endpoints of the study were variations in systemic metabolism, insulin, and neurotrophic signaling, neuroinflammation, changes in levels of Aβ-and tau-pathology or cognitive performances. Age-matched 3xTg-AD animals fed with a standard diet or undergoing HFD and treated with vehicle were used as controls.

## Materials and Methods

### Animals and treatment paradigm

All the procedures involving the animals and their care were approved by the Local Institutional Ethics Committee (Comitato Etico Interistituzionale per la Sperimentazione Animale [CEISA] protocol no. 17; Min. IDD: DGSAF/14264). Animal handling was performed in compliance with national and international laws and policies. All efforts were employed to reduce the number of animals and their suffering upon all the experimental procedures. 3xTg-AD mice [B6;129-Tg(APPSwe,tauP301L)1Lfa *Psen1^tm1Mpm^*/Mmjax] were purchased from Jackson Laboratory, bred in the Center of Excellence on Aging and Translational Medicine (CeSI-MeT) animal facility, housed on 12-12 hours light/dark cycle, and provided with, until treatment allocation, *ad libitum* access to standard chow and water. A total of forty-six 3xTg-AD mice (21 males and 25 females) were enrolled at six months of age (m.o.a.) and randomly assigned to a 6-month control or high fat dietary regimen. 3-month after the beginning of the dietary treatment, mice of the control (3xTg-AD^CD^) and high fat (3xTg-AD^HFD^) groups were randomly assigned to a 3-month administration of exenatide or vehicle (PBS). HFD was purchased from Altromin. In this obesity-inducing chow, 60% of the energy derives from fats. CD consists of a standard chow with 13% of energy deriving from fats. Exenatide or vehicle administration were performed as previously described ^26^. Briefly, exenatide (500 μg/kg body weight) or vehicle were administered via intraperitoneal injection five days per week. Treatments were also maintained during the behavioral testing phase. The exenatide lyophilized powder was provided by Eli Lilly.

### Insulin sensitivity and glucose tolerance tests

Insulin sensitivity and glucose tolerance were assessed at 6, 9, and 12 m.o.a. by employing the intraperitoneal insulin tolerance test (ITT) and the glucose tolerance test (GTT), respectively. For ITT, after a six-hour fasting period, mice were injected with 0.75 unit/kg of human insulin (Sigma-Aldrich). For, GTT, after an overnight fasting period (≈ 16 h), mice were injected with 1g/kg glucose (Sigma-Aldrich). Glycemia was measured from vein tail blood drop with a Freestyle InsuLinx glucometer (Abbott). Measurements were performed 5, 15 30, 45, 60, 120, and 180 min after the insulin or glucose administration. Insulin-related measurements were halted at the 30^th^ min to avoid hypoglycemia.

### Plasma insulin assay and HOMA-IR assessment

Plasma insulin concentrations were determined with the Ultrasensitive Insulin ELISA kit (Mercodia) following the manufacturer instructions. Fasting glucose and insulin concentrations for the Homeostasis Model Assessment of Insulin Resistance (HOMA-IR) calculations were employed as [insulin (pM/L) X glucose (mM/L) / 22.5].

### Tissue collection

At the end of treatments and following behavioral tests and metabolic analyses, mice were anesthetized, killed, and tissue samples harvested for biochemical analysis. Brains were halved into 2 hemispheres. For immunohistochemical (IHC) analyses, one hemisphere was collected in a Carnoy solution, kept for 2 days at 4° C, ethanol-washed, and paraffin-embedded until sectioning. For Western blot (WB) analysis, each hemisphere was dissected into subregions (hippocampus, whole cortex, and cerebellum), snap-frozen in liquid nitrogen, and stored at −80 °C until sampling.

### Aβ and p-tau immunohistochemistry

Five µm sections of Carnoy-fixed and paraffin-embedded brains of 3xTg-AD mice from the four experimental groups were stained using purified mouse monoclonal antibodies raised against human Aβ (clone DE2B4, 1:200 dilution, overnight incubation, Abcam) and p-tau (Thr_231_; AT180, Pierce Protein Research Products) as previously reported ^25^. Briefly, antigen retrieval was performed in 10 mmol/l sodium citrate buffer (pH 6.0) by a thermostatic bath at 100 °C for 10 min for Aβ and by microwave treatment at 750 W for 10 min for p-tau. The anti-mouse EnVision kit (Agilent) was used for signal amplification. In control sections, the specific primary antibodies were replaced with isotype-matched immunoglobulins. In the hippocampus, Aβ deposition was quantified by counting the number of Aβ plaques using a 10x magnification. The immunostaining signal evaluation for p-tau was performed by counting stained pixels on hippocampal neurons using Photoshop (Adobe Systems) as previously reported ^25^.

### Western blot analysis

Brain regions were lysed in ice-cold RIPA buffer containing [in %]: 0.5 sodium deoxycholate, 1 Nonidet P-40, 0.1 SDS, 1 protease and phosphatase inhibitor cocktails, and 5 mM EDTA, pH 7.4. Protein lysates (10 μg) were separated on a SDS-polyacrylamide gel (9%–13% gradient) and blotted onto PVDF membrane. Nonspecific binding sites were blocked with 5% non-fat dry milk (Bio-Rad Laboratories) in Tris-buffered saline (TBS) containing [in mM]: 20 Tris–HCl, 150 NaCl, pH 7.4, for 30 minutes at room temperature. Membranes were incubated overnight at 4 °C with the primary antibodies, diluted with TBS supplemented with 0.1% Tween 20 (TBS-T) and 5% nonfat dry milk. A list of the WB employed antibodies and their dilution is available in the Supplementary Table S1. Peroxidase-conjugated secondary anti-rabbit or anti-mouse IgG antibodies (1:10,000; Vector Laboratories) were used. Chemiluminescent signals were visualized by enhanced chemiluminescence (EuroClone), following the manufacturer instructions. Relative densities were determined and normalized to a housekeeping protein (actin) using ImageJ software. Values are given as relative units or phosphoprotein/total protein ratio, and calculated as: (phosphoprotein/loading control)/(total protein/loading control).

### Morris water maze test

The Morris Water Maze (MWM) test was performed as previously described ^30^. Briefly, the MWM apparatus (Panlab) consists in a circular pool (1.2 m diameter) filled with warm water (22 ± 1 °C). The pool is placed in a noise-isolated room containing several intra-and extra-maze visual cues. Mice were trained to swim in the pool and reach a circular platform located 2 cm beneath the water surface. Mice that failed to find the platform within 90 seconds were manually guided to it and allowed to remain there for 10 seconds. Mice performed four trials per day for four consecutive days. Spatial memory performances were assessed 1.5 and 24 hours after the end of the last training trial. The probe test consisted of a 60-second free swim in the pool in which the platform has been removed. Probe tests were recorded for subsequent analysis with the Smart tracking software (Panlab). Performances were evaluated in terms of time employed to reach the location where the platform used to be (escape latency), number of crosses over the platform location, and time spent in the target (T target) quadrant.

### The novel object recognition test

The Novel Object Recognition (NOR) test was performed as previously described ^31^. Briefly, in the habituation phase, 3xTg-AD mice were placed for 10 min per day for 2 consecutive days in an empty cage. On day 3, mice were placed in a cage containing two identical objects spaced ∼15 cm apart and allowed to explore the objects for 8 min. At the end of each trial, objects were thoroughly cleaned with ethanol and air dried. On day 4, mice were placed in the experimental cage containing the objects that have been previously presented. One of these objects was left in the previous location (familiar location) while the other one was placed in a new position (novel location). Mice were then allowed to explore the two objects for 5 min (probe test). The test was videotaped for subsequent analysis that was manually performed in blind conditions. Scoring of the NOR performance was analyzed in terms of time spent to explore both objects, presented in a familiar location and novel one. Mice were considered explorative when the head was within 1 cm from the object with the neck extended and vibrissae moving. Proximity or chewing did not count as exploration. Analyzed parameters were the percentage of time spent with the object in the novel location and the discrimination index (DI). DI was calculated as follows: ((A–B)/(A+B)) × 100, where A is the time spent to explore the object in the novel location and B is the time spent on the object in the familiar one ^31^.

### Data analysis

No statistical methods were used to predetermine sample size. Statistical analysis was performed by one-way ANOVA followed by Tukey’s post-hoc test. For IHC data Kruskal-Wallis test followed by Tamhane post-hoc test was performed. SPSS Version 15.0 was used throughout. By conventional criteria, the level of significance was set at p < 0.05. Data were expressed as mean ± standard error of the mean unless otherwise indicated.

## Results

### Exenatide treatment has no effect on body weight and glucose metabolism in 3xTg-AD^CD^ and 3xTg-AD^HFD^ mice

Male and female 3xTg-AD mice (n=46) at 6 m.o.a. were randomly assigned to a control (3xTg-AD^CD^; n=24) or HFD (3xTg-AD^HFD^; n=22) for 6 months. At 9 m.o.a., 3xTg-AD^CD^ and 3xTg-AD^HFD^ mice were subjected to intraperitoneal injections of exenatide (500 µg/kg, five days per week; n=11 and 9, respectively) or vehicle (saline; n=13 for both groups). Compared to 3xTg-AD^CD^ animals, 3xTg-AD^HFD^ mice exhibited a significant increase in body weight (Fig. 1b; p<0.001). Unexpectedly, no differences in body weight were found when comparing vehicle-and exenatide-treated animals (Fig. 1b; p>0.05).

**Figure 1.**
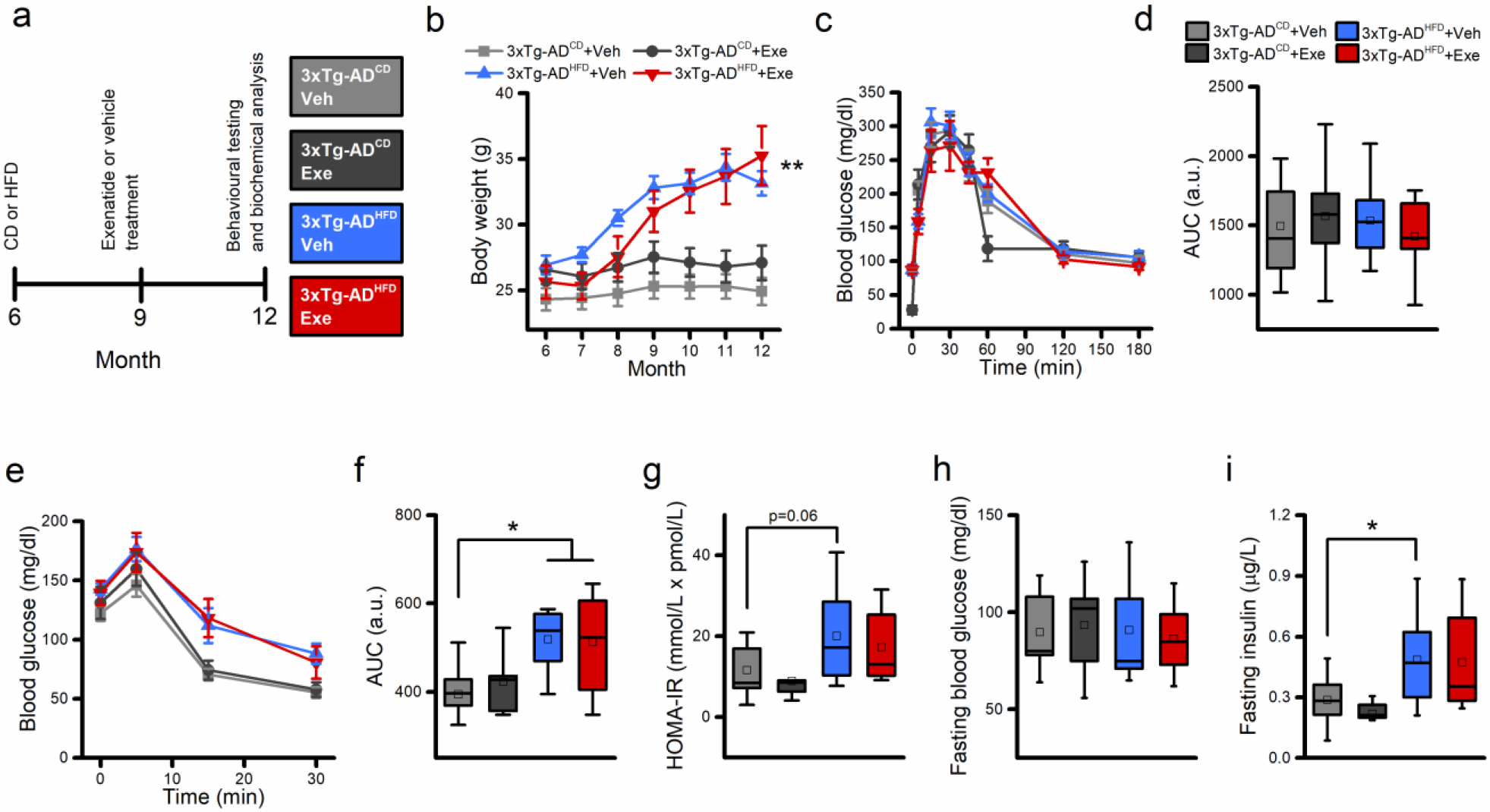
Effects of HFD and exenatide on systemic metabolism. (a) The pictogram illustrates the experimental paradigm employed in the study. (b) Graphs depict body weight changes of 3xTg-AD^CD^ (Veh, n=13; Exe, n=11) or 3xTg-AD^HFD^ (Veh, n=13; Exe, n=9) mice treated with either vehicle or exenatide. (c) Graphs depict the intra-peritoneal GTT (glucose tolerance test) curve of 12 m.o. vehicle- and exenatide-treated 3xTg-AD^CD^ or 3xTg-AD^HFD^ mice. (d) Box charts illustrate the GTT quantifications expressed as area under the curve (AUC). (e) Graphs illustrate the intra-peritoneal ITT (insulin tolerance test) of the four study groups. (f) Box charts illustrate the ITT quantifications expressed as area under the curve (AUC). (g) Box charts indicate changes of the Homeostasis Model Assessment of Insulin Resistance (HOMA-IR) scores in the four groups. (h-i) Box charts illustrate changes in fasting blood glucose and insulin levels. Data are expressed as mean ± SEM or as box charts where center lines show median, center boxes show mean, box limits indicate 25^th^ and 75^th^ percentiles, and whiskers extend 1.5 times the interquartile range. Means were compared by one-way ANOVA followed by Tukey post-hoc test. * indicates p<0.05; ** indicates p<0.01. Abbreviations: Veh, vehicle; Exe, exenatide.

HFD promotes IR and T2DM ^32^. To evaluate the HFD-dependent metabolic changes, glucose metabolism and insulin sensitivity were measured in vehicle-and exenatide-treated 3xTg-AD^CD^ and 3xTg-AD^HFD^ mice. Mice were sampled at 12 m.o.a. No metabolic differences were observed at 6 or 9 m.o. among the study groups (Supplementary fig. 1). Analysis of the intra-peritoneal GTT, a measure of the body ability to metabolize glucose, showed no diet-or drug-related effects in the study groups (Fig. 1c-d; p>0.05). Similarly, no differences were observed in fasting blood glucose levels (Fig. 1h; p>0.05). Evaluation of the ITT, to assess the body response to the exogenous administration of insulin, showed that 3xTg-AD^HFD^ mice exhibit altered ITT when compared to 3xTg-AD^CD^ animals (Fig. 1e-f; p<0.01). In addition, compared to 3xTg-AD^CD^ animals, 3xTg-AD^HFD^ mice showed a mild increase in fasting plasma insulin levels (Fig. 1i; p=0.06). These insulin-related parameters were not affected by the use of exenatide in both diets (Fig. 1e-i; p>0.05), thereby indicating a lack of metabolic effect of the compound.

The HOMA-IR is a mathematical formula that takes into account resting levels of glucose and insulin and provides an indirect estimation of IR. Compared to 3xTg-AD^CD^ animals, the analysis of this parameter showed a trend toward increased HOMA-IR levels in 3xTg-AD^HFD^ mice (Fig. 1g; p=0.06). No exenatide-driven effects were observed in 3xTg-AD^HFD^ or 3xTg-AD^CD^ mice (Fig. 1g; p<0.05).

### Aβ-and tau-pathology in 3xTg-AD^CD^ and 3xTg-AD^HFD^ mice

Previous studies have shown that the HFD administration exacerbates the development of Aβ and tau pathology in preclinical AD models including the 3xTg-AD mice ^33,34^. The IHC analysis of hippocampal Aβ plaque loads and p-tau immunoreactivity of the study animals revealed no differences when comparing 3xTg-AD^CD^ and 3xTg-AD^HFD^ mice (Fig. 2a-d). In agreement with previous findings ^25^, exenatide treatment did not affect Aβ and tau levels in 3xTg-AD^CD^ and 3xTg-AD^HFD^ animals (Fig. 2a-d).

**Figure 2.**
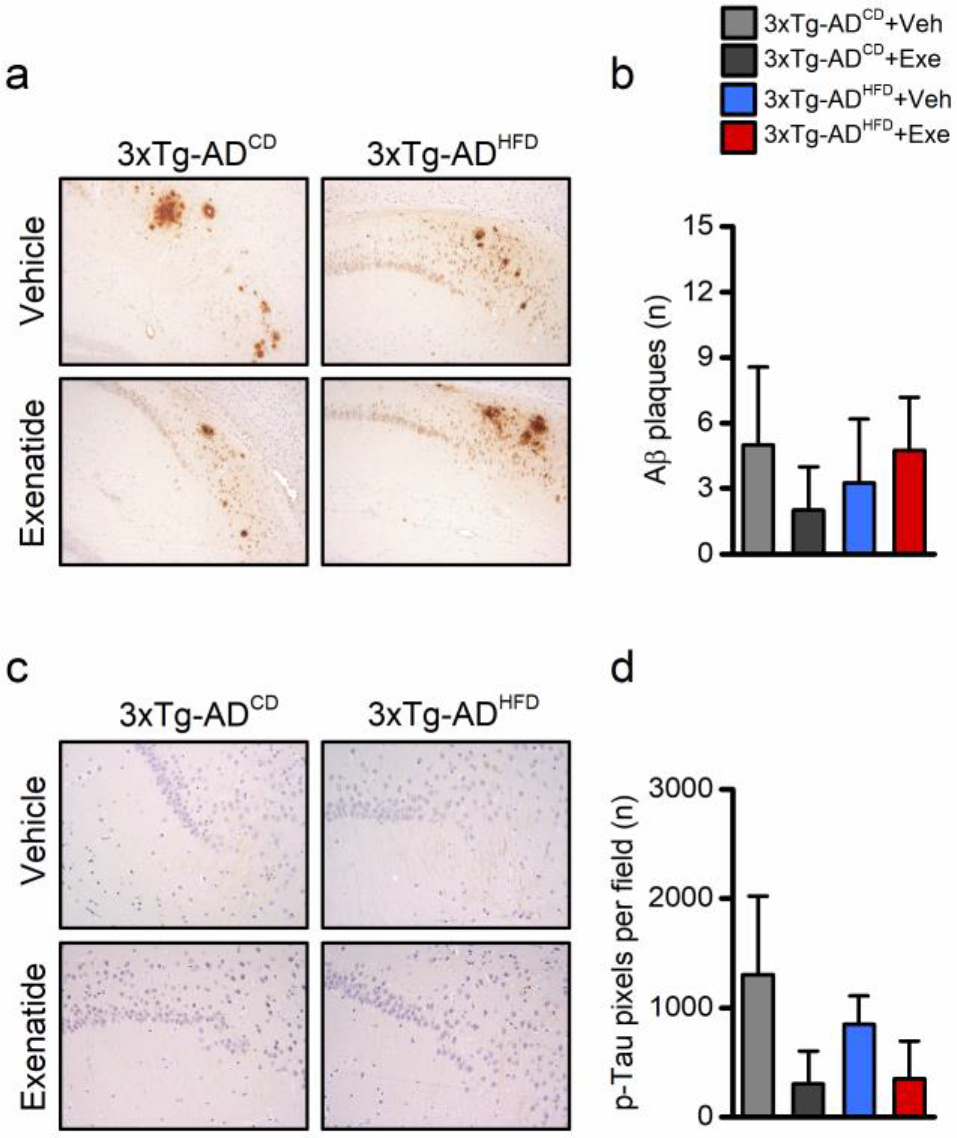
Effects of HFD and exenatide on Aβ and tau pathology. (a) Representative images of Aβplaquedeposition in the hippocampal CA1 region of 3xTg-AD^CD^ (Veh, n=5; Exe, n=3) or 3xTg-AD^HFD^ (Veh, n=4; Exe, n=4) mice treated with either vehicle or exenatide. (b) Bar graph depicts Aβ deposit quantification expressed as number of plaques per field. (c) Representative images of intraneuronal p-tau immunoreactivity in the hippocampal CA1 region of 3xTg-AD^CD^ (Veh, n=5; Exe, n=3) or 3xTg-AD^HFD^ (Veh, n=4; Exe, n=3) mice treated with either vehicle or exenatide. Data were analyzed by Kruskal-Wallis test followed by Tamhane post-hoc test. No statistically significant differences were observed among the four study groups. Abbreviations: Veh, vehicle; Exe, exenatide.

### Exenatide treatment positively affects BDNF signaling in 3xTg-AD^CD^ mice and prevents the development of neurotrophic signaling impairment in 3xTg-AD^HFD^ animals

HFD administration impairs BDNF neurotrophic signaling along with the downstream activation of the cAMP response element–binding (CREB) protein, a transcription factor essential for synaptic plasticity and learning and memory processes ^35,36^. Compared to 3xTg-AD^CD^ animals, WB analysis performed on hippocampal, cortical, and cerebellar homogenates obtained from the study animals showed decreased BDNF levels in 3xTg-AD^HFD^ mice (Fig. 3a; p<0.01). This result was mirrored by an HFD-driven hippocampal reduction of the active forms of key plasticity related modulators. Reduced levels of phosphorylated TrkB (pTrkB; the high-affinity BDNF receptor), phosphorylated ERK5 (a downstream effector of the BDNF/TrkB axis), and phosphorylated CREB (pCREB; Figs. 3b-d; p<0.01) were observed in 3xTg-AD^HFD^ mice. As BDNF signaling is a critical regulator of structural plasticity, we investigated the HFD-related effects on phosphorylated Synapsin I (pSyn) and PSD95, two proteins known to be involved in synapse stabilization at pre-and post-synaptic levels ^37,38^. In line with the BDNF results, 3xTg-AD^HFD^ mice showed a marked decrease of the two proteins when compared to 3xTg-AD^CD^ animals (Fig. 3e-f; p<0.001).

**Figure 3.**
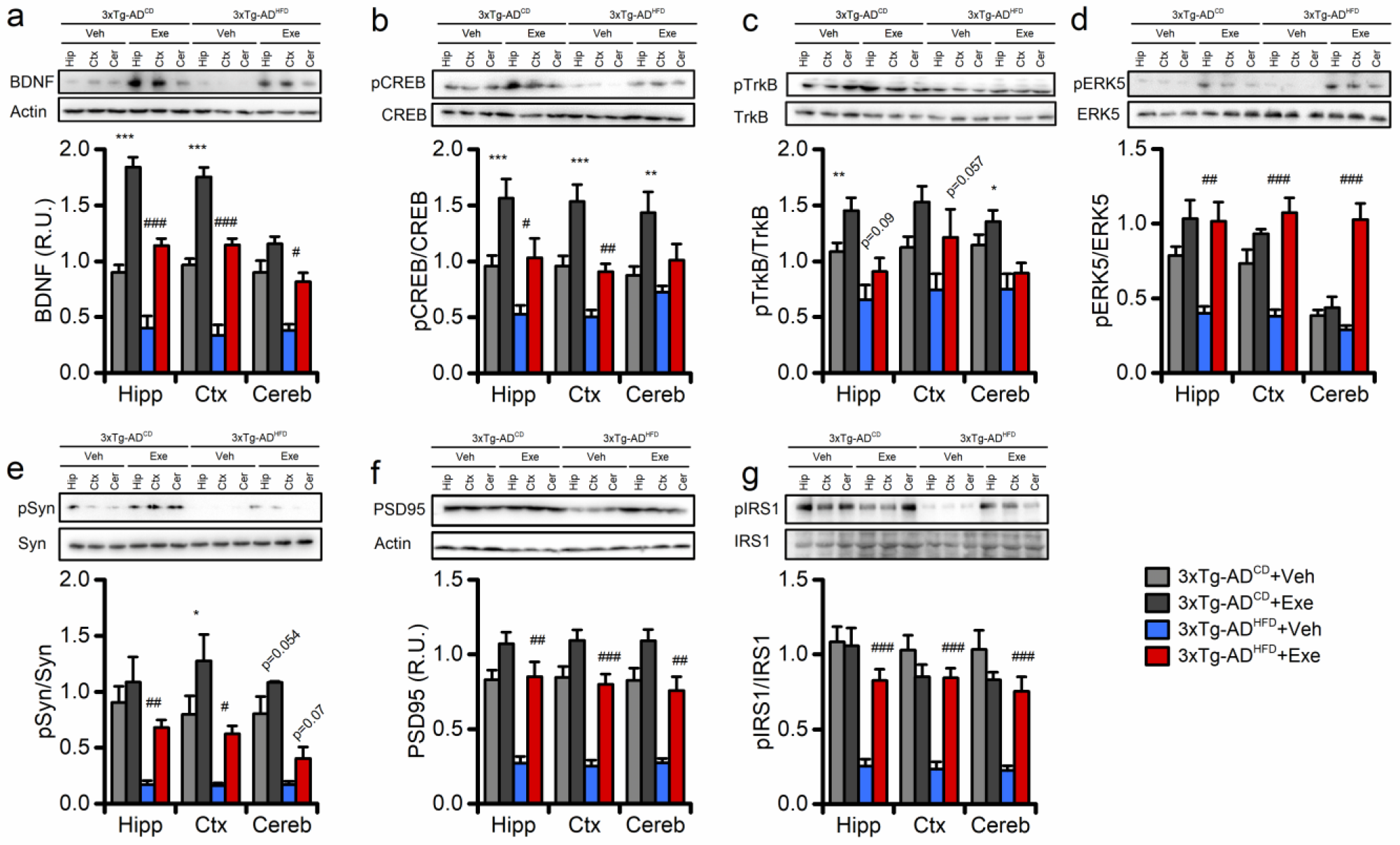
Effects of HFD and exenatide on BDNF neurotrophic signaling. Western blots show exenatide-or vehicle-driven effects on BDNF-related signaling in the hippocampus (Hip), cortex (Ctx), and cerebellum (Cer) of 12 m.o. 3xTg-AD^CD^ and 3xTg-AD^HFD^ mice; images are representative of 3-5 independent experiments. (a) Bar graphs depict BDNF levels in the four study groups (n=5). (b) Bar graphs depict levels of pCREB in the four study groups (n=5). (c) Bar graphs depict pTrkB levels in the four study groups (n=5). (d) Bar graphs depict pERK5 levels in the four study groups (n=5). (e) Bar graph depicts levels of pSyn in the four study groups (n=3). (f) Bar graphs depict expression levels of PSD95 in the four study groups (n=5). Data show mean ± SEM of relative units (R.U.). Means were compared by one-way ANOVA followed by Tukey post-hoc test. * indicates p<0.05, ** p<0.01, *** p<0.001 of 3xTg-AD^CD^ + Veh versus 3xTg-AD^CD^ + Exe; # indicates p<0.05, ## p<0.01, ### p<0.001 of 3xTg-AD^HFD^ + Veh versus 3xTg-AD^HFD^ + Exe. Abbreviations: Veh, vehicle; Exe, exenatide.

Compared to vehicle-treated animals, exenatide-treated mice showed a significant increase in BDNF, pERK5, pCREB, pSyn, and PSD95 (Fig. 3a-f). The drug also promoted increased pTrkB levels (Fig. 3c). These changes showed a trend toward statistical significance when analyzed with ANOVA (Tukey post-hoc test; p=0.09) and were statistically significant when not corrected for multiple comparisons (Student t-test; p=0.02). In the 3xTg-AD^HFD^ mice, exenatide prevented the development of HFD-induced impairment of BDNF signaling (Fig. 3a). Exenatide also promoted enhanced BDNF signaling in 3xTg-AD^CD^ mice, thereby suggesting that the molecule produces a neurotrophic drive independently of the dietary regimen (Fig. 3a; p<0.001).

As GLP-1R signaling modulates insulin-related pathways, we evaluated the insulin receptor substrate 1 (IRS-1) phosphorylation status at serine 1101, a modification known to inhibit downstream insulin signaling ^39^. WB analysis of this parameter showed decreased pIRS-1 levels in 3xTg-AD^HFD^ mice (p<0.001), a change that was reverted by exenatide administration (Fig. 3g). No drug-related effects were observed in 3xTg-AD^CD^ animals (Fig. 3g; p>0.05).

### Exenatide treatment decreases p75NTR activation in 3xTg-AD^CD^ mice and prevents the neurotoxic signaling in 3xTg-AD^HFD^ animals

The biologically mature, and thus active, form of BDNF originates from the proteolytic cleavage of proBDNF. In contrast with the plasticity effects of BDNF, proBDNF, acting on the high-affinity P75 neurotrophin receptor (P75NTR), activates apoptotic signaling ^40^. WB analysis of hippocampal, cortical, and cerebellar lysates of the study groups showed increased proBDNF levels in 3xTg-AD^HFD^ mice (Fig. 4a; p<0.01). HFD promoted p75NTR overexpression and phosphorylation as well as activation of JNK (pJNK) and ERK_1,2_ (pERK_1,2_), two downstream effectors of the proBDNF/p75NTR signaling cascade (Fig. 4b-d; p<0.01). Compared to vehicle-treated 3xTg-AD^CD^ or 3xTg-AD^HFD^ mice, exenatide administration significantly reduced proBDNF signaling in 3xTg-AD^CD^ or 3xTg-AD^HFD^ animals (Fig. 4a). In 3xTg-AD^HFD^ mice, the molecule was found to revert the HFD-induced activation of proBDNF/p75NTR signaling (Fig. 4b-d).

**Figure 4.**
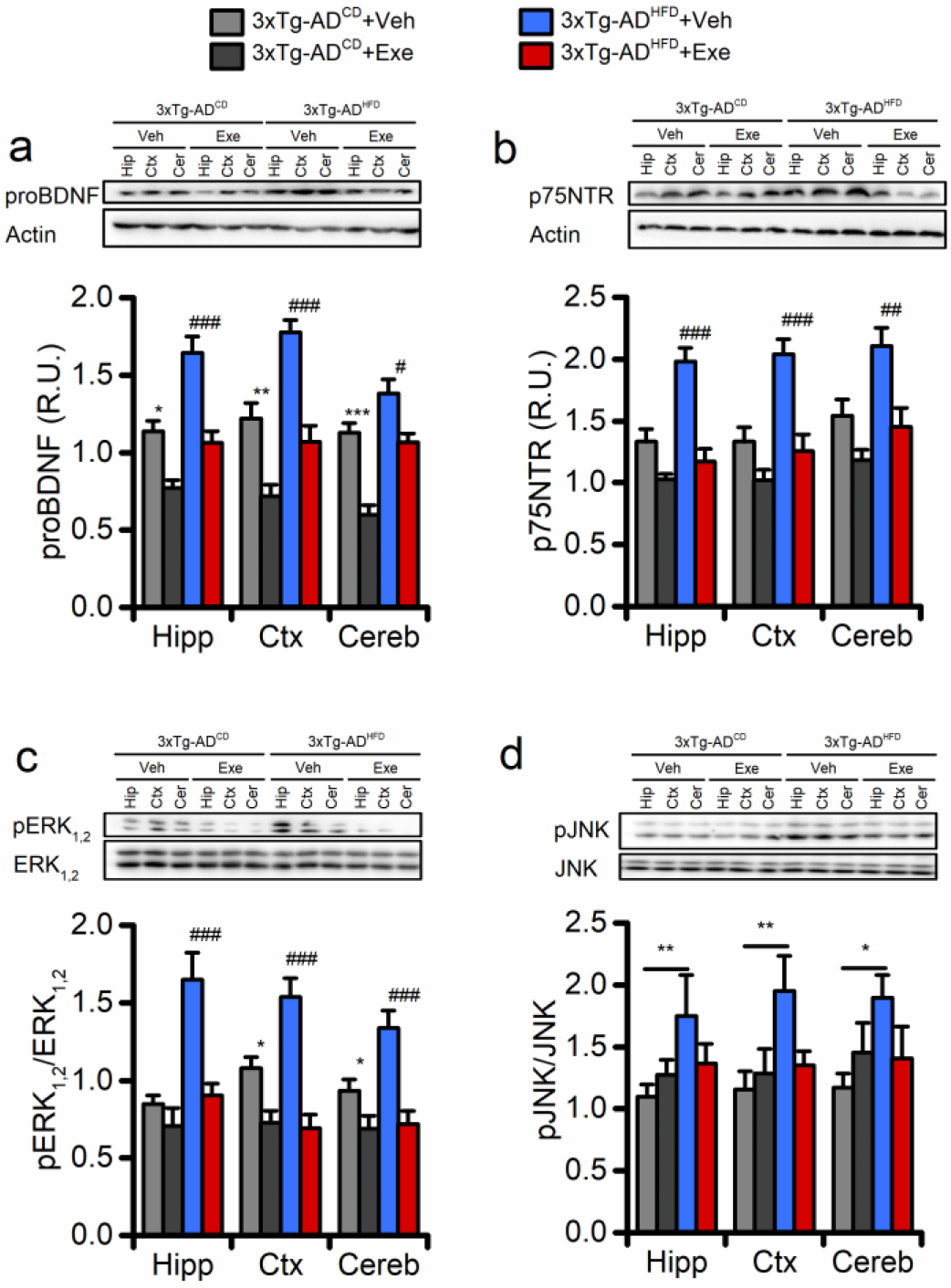
Effects of HFD and exenatide on proBDNF neurotoxic signaling. Western blots show exenatide-or vehicle-driven effects on proBDNF-related signaling in the hippocampus (Hip), the cortex (Ctx), and the cerebellum (Cer) of 12 m.o. 3xTg-AD^CD^ and 3xTg-AD^HFD^ mice; images are representative of 5 independent experiments. (a) Bar graphs depict proBDNF levels in the four study groups (n=5). (b) Bar graphs depict expression levels of p75NTR in the four study groups (n=5). (c) Bar graphs depict pERK_1,2_ levels in the four study groups (n=5). (d) Bar graphs depict pJNK levels in the four study groups (n=5). Data show mean ± SEM of relative units (R.U.). Means were compared by one-way ANOVA followed by Tukey post-hoc test. * indicates p<0.05, ** p<0.01, *** p<0.001 of 3xTg-AD^CD^ + Veh versus 3xTg-AD^CD^ + Exe; # indicates p<0.05, ## p<0.01, ### p<0.001 of 3xTg-AD^HFD^ + Veh versus 3xTg-AD^HFD^ + Exe. Abbreviations: Veh, vehicle; Exe, exenatide.

### Exenatide treatment modulates the inflammatory response in 3xTg-AD^HFD^ mice

The HFD is known to promote a pro-inflammatory state through the activation of lipid-mediated signaling ^41^. A growing body of evidence indicate that neuroinflammation is present in AD ^42^. To address the role of inflammation in our experimental setting, we performed a WB analysis of the pro-inflammatory mediator NF-κB. We also evaluated PPARs, the anti-inflammatory peroxisome proliferator-activated receptor proteins ^43,44^. A significant increase in NF-κB levels was found in 3xTg-AD^HFD^ mice, a phenomenon reverted by the exenatide administration (Fig. 5a; p<0.01). No exenatide-driven effects were observed in 3xTg-AD^CD^ animals (Fig. 5a; p>0.05). In 3xTg-AD^HFD^ mice, the analysis of the diet-related effects on PPARs revealed a net increase in levels of the PPARα and PPARγ isoforms (Fig. 5b-d; p<0.001). The effects were inhibited by the exenatide administration (Fig. 5b-d). No effects were observed in exenatide-treated 3xTg-AD^CD^ mice (Fig. 5b-d; p>0.05). In addition, no diet- or drug-related effects were observed when analyzing PPARβ/δ levels (Fig. 5c).

**Figure 5.**
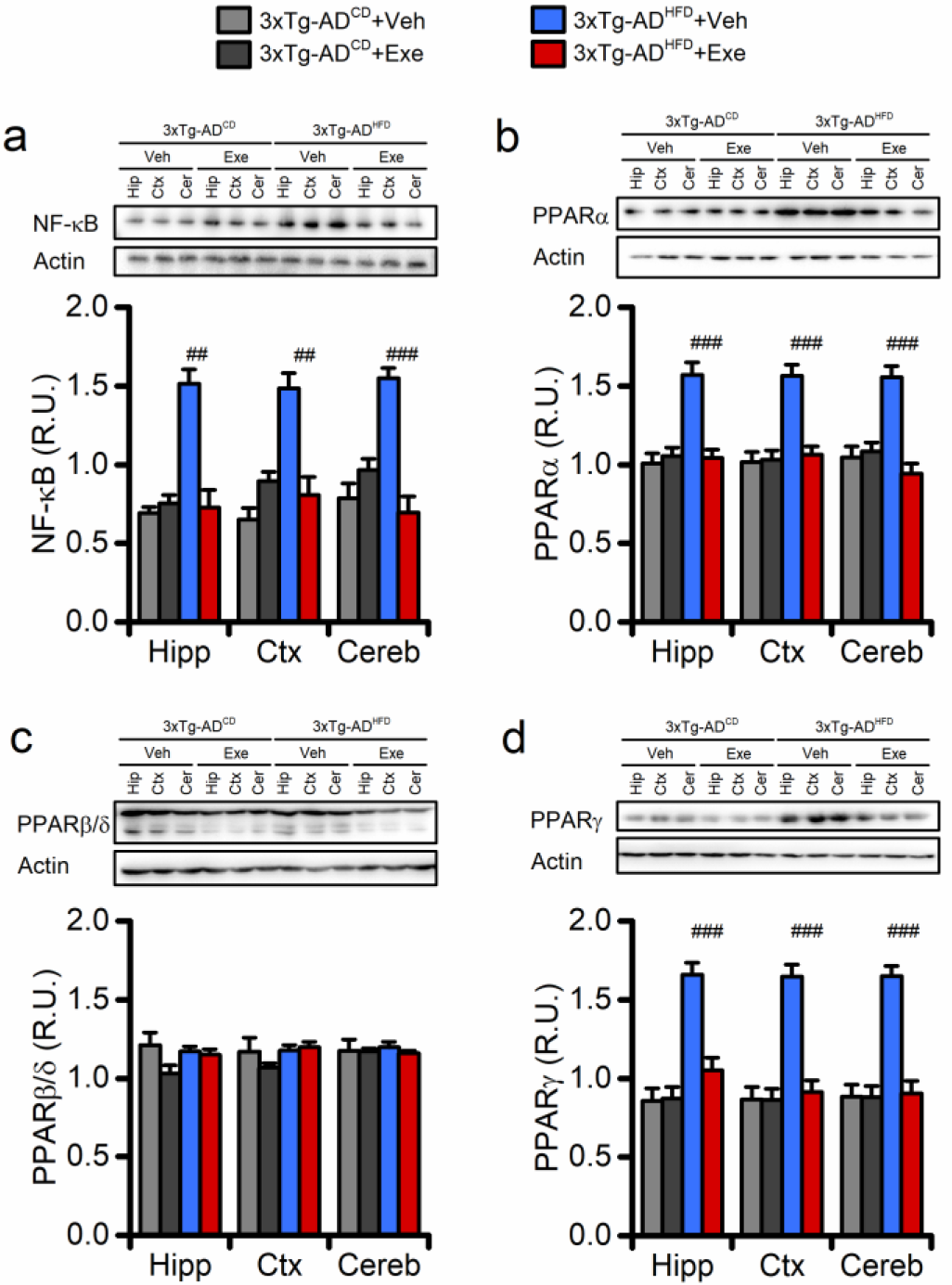
Effects of HFD and exenatide on brain inflammation. Western blots show exenatide-or vehicle-driven effects on pro-and anti-inflammatory markers in the hippocampus (Hip), the cortex (Ctx), and the cerebellum (Cer) of 12 m.o. 3xTg-AD^CD^ and 3xTg-AD^HFD^ mice; images are representative of 3-5 independent experiments. (a) Bar graphs depict expression levels of NF-κB in the four study groups (n=3). (b) Bar graphs depict expression levels of PPARα in the four study groups (n=5). (c) Bar graphs depict expression levels of PPARβ/δ in the four study groups (n=5). (d) Bar graphs depict expression levels of PPARγ in the four study groups (n=5). Data show mean ± SEM of relative units (R.U.). Means were compared by one-way ANOVA followed by Tukey post-hoc test. ## indicates p<0.01, ### p<0.001 of 3xTg-AD^HFD^ + Veh versus 3xTg-AD^HFD^ + Exe. Abbreviations: Veh, vehicle; Exe, exenatide.

### Exenatide treatment has no effects on learning and memory performances in 3xTg-AD^CD^ and 3xTg-AD^HFD^ mice

Cognitive effects of HFD and exenatide administration were evaluated in the study groups. To that aim, we employed the MWM test, an experimental setting that evaluates hippocampus-dependent spatial memory ^45^. Unexpectedly, compared to 3xTg-AD^CD^ animals, 3xTg-AD^HFD^ mice did not show deficits in learning or long-term memory (Fig. 6a-e; p>0.05). Exenatide treatment resulted in no effects on cognition (Fig. 6a-e; p>0.05). Cognitive performances were also evaluated with the NOR test. The NOR test investigates the hippocampus-dependent spatial memory performances. Compared to the MWM, the NOR is considered more sensitive as the test is less affected by stress-related biases ^31^. Compared to 3xTg-AD^CD^ animals, 3xTg-AD^HFD^ mice did not show differences in the time spent with either the object in the novel location or the DI (Fig. 6f-h; p>0.05). Exenatide administration had no effects on these parameters (Fig. 6f-h; p>0.05).

**Figure 6.**
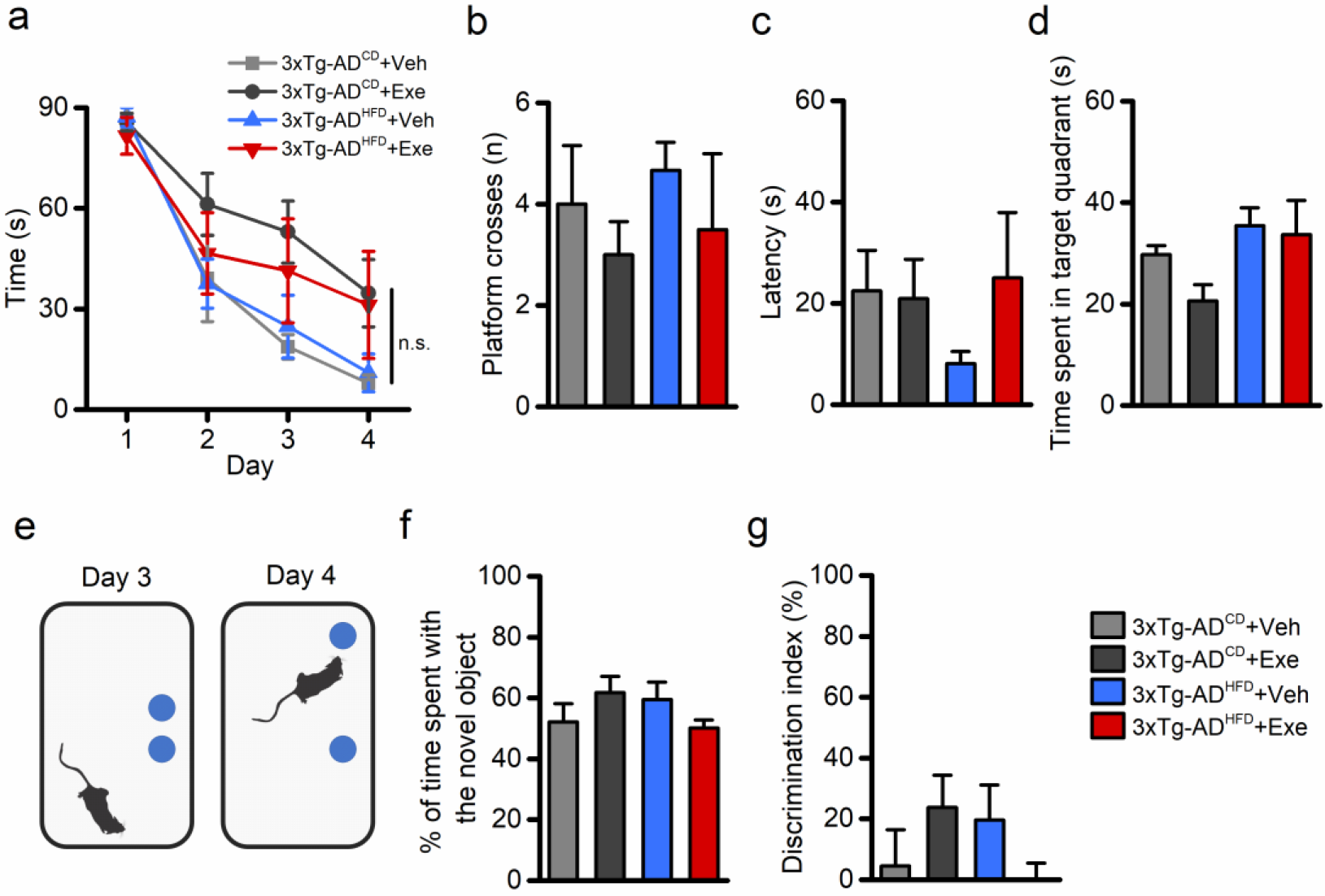
Effects of HFD and exenatide on memory performances. Memory performances were evaluated, in the four study groups, with the Morris Water Maze (MWM) and with the Novel Object Recognition (NOR) tests. (a) The graph depicts the learning curve of 3xTg-AD^CD^ + Veh (n=6), 3xTg-AD^CD^ + Exe (n=7), 3xTg-AD^HFD^ + Veh (n=6), and 3xTg-AD^HFD^ + Exe (n=5) mice as assessed upon the 4-day training sessions. Analysis of MWM data revealed no statistically significant differences in learning performances between the four groups (p>0.05). (b) Bar graphs depict the number of crosses (the number of times each mouse crosses the location where the platform used to be) in the four study groups. (c) Bar graphs depict the latency (the time spent to reach the location where the platform used to be) in the four study groups. (d) Bar graphs depict the time spent in the target quadrant (the quadrant where the platform used to be) in the four study groups. (e) The pictogram illustrates the NOR experimental paradigm employed for testing 3xTg-AD^CD^ + Veh (n=12), 3xTg-AD^CD^ + Exe (n=11), 3xTg-AD^HFD^ + Veh (n=12), and 3xTg-AD^HFD^ + Exe (n=9). (f) Bar graphs depict the percentage of time spent with the object moved to a novel location in the four study groups. (g) Bar graphs depict the discrimination index (DI; the difference between time spent exploring novel and familiar objects) in the four study groups.

## Discussion

In this study, we found that, in 3xTg-AD^HFD^ mice, 3-month treatment with exenatide enhances BDNF signaling and reduces inflammation. Positive neurotrophic effects were also observed in exenatide-treated 3xTg-AD^CD^ animals. These findings are in line with previous results supporting a role for GLP-1R agonism in the beneficial modulation of neurotrophic signaling ^26,28,46^. Results of the study also revealed distinct molecular effects exerted by HFD on the AD-like background of the 3xTg-AD animals.

### Molecular effects of HFD on 3xTg-AD mice

HFD administration promoted weight increases and significant IR alterations but did not affect the levels of blood glucose of the mice (Fig. 1). In line with previous findings, compared to 3xTg-AD^CD^ animals, 3xTg-AD^HFD^ mice developed signs of IR ^47^. However, the HFD regimen did not induce a chronic hyperglycemic state or diabetes in the 3xTg-AD animals (Fig. 1h). This result may be explained by the duration of our dietary regimen ^48^ as longer (> 6 months) HFD is required to produce T2DM.

Previous findings have shown that HFD affects cognition by impairing neurotrophic signaling ^35^. In agreement with this notion, in 3xTg-AD mice, HFD promoted a significant decrease in the activation of CREB and BDNF/TrkB signaling (Fig. 3a-c). In our AD model, HFD also decreased levels of pERK5, a BDNF-activated kinase involved in neuronal survival ^49,50^, PSD95, a postsynaptic density marker associated with BDNF-driven structural plasticity ^37^, and pSyn, a BDNF-activated protein known to promote synaptic vesicle clustering and neurotransmitter release ^38^.

In the CNS, the trophic effects of neurotrophins are counteracted by the activity of pro-neurotrophins, their immature precursor forms. Mature neurotrophins originate from the proteolytic cleavage of pro-neurotrophins, and changes in the balance of mature/immature forms affect neuronal functioning. proBDNF, the precursor form of BDNF, exhibits a high binding affinity for p75NTR ^51^. p75NTR, once activated, promotes pro-apoptotic pathways through ERK_1,2_- and JNK-dependent signaling. In 3xTg-AD^HFD^ mice, we found increased proBDNF levels along with the activation of p75NTR, ERK_1,2_, and JNK (Fig. 4). These results are in line with previous studies showing that HFD and/or IR promote JNK signaling ^16,32^. Our results provide a potential causal link between HFD, the activation of proBDNF signaling, and JNK. The HFD-driven activation of the proBDNF/p75NTR axis was paralleled by a net reduction in levels of pre- and post-synaptic markers like pSyn and PSD95 (Fig. 3e-f). These findings are in agreement with recent evidence indicating that proBDNF-related signaling, occurring via p75NTR, negatively affects neuronal functioning and synaptic remodeling ^52^. Our findings also identify, a negative role for ERK_1,2_. Although ERK_1,2_ activation has been previously considered as a critical step for the BDNF signaling cascade, a growing body of recent evidence has challenged the positive effect of pERK_1,2_ and shown that the molecules participate in several death-related mechanisms (reviewed in ^53^). In that regard and in line with our previous findings ^26^, our results support the idea that ERK_1,2_ promotes divergent and bimodal effects and indicate a deleterious HFD-driven activation of ERK_1,2_ ^54^.

Inflammation plays a central role in the pathogenesis of neurodegenerative conditions, including AD ^42^. Our findings support the idea that the HFD is a potent trigger of NF-κB, a transcription factor involved in the expression of inflammatory-related cytokines and chemokines (Fig. 5a) ^55,56^. In our study, HFD was also found to increase the expression of PPARα and PPARγ (but not PPARβ/δ), two proteins known to counteract pro-inflammatory pathways (Fig. 5b-d) ^43^. This finding may be viewed as a compensatory mechanism set in motion to counteract the HFD-driven inflammatory response ^44^. This hypothesis is supported by recent findings showing that, upon active brain inflammation, PPARα and PPARγ but not PPARβ/δ are selectively expressed in microglia ^57^ (Fig. 5b-d).

The HFD did not affect Aβ pathology in our 3xTg-AD mice. This finding diverges from previous reports showing that fat-enriched diets can exacerbate the amyloid pathology ^33,34^ but are in line with other studies reporting the lack of effect of HFD on Aβ plaque deposition ^54,58^ (Fig. 2a-b). No changes were also observed in p-tau levels when comparing 3xTg-AD^CD^ and 3xTg-AD^HFD^ mice (Fig. 2c-d).

### Molecular effects of exenatide on 3xTg-AD^CD^ and 3xTg-AD^HFD^ mice

Exenatide was not effective in restoring the insulin sensitivity in the 3xTg-AD^HFD^ mice, a lack of efficacy likely because the compound promotes regulatory activities only in the presence of patent signs of hyperglycemia or diabetes ^26,59^. 3xTg-AD mice at 12 m.o.a. are devoid of metabolic deficits ^31^, thereby not offering the pathological background on which exenatide can work.

Previous *in vitro* and *in vivo* findings indicate that the GLP1-R agonism exerted by endogenous or exogenous synthetic ligands activates CREB and upregulates BDNF levels ^60,61^. We have recently shown that exenatide, administered to adult wild-type mice positively modulates the BDNF-TrkB neurotrophic axis ^26^. In this study, exenatide was found to increase the BDNF-related signaling in the hippocampus of 3xTg-AD^CD^ mice and partially restored it to control levels in 3xTg-AD^HFD^ animals (Fig. 3a-c). Given the central role played by BDNF in structural plasticity, it is conceivable that some of the exenatide effects may have worked on synaptic targets. In agreement with this hypothesis, the exenatide-related potentiation of BDNF signaling was paralleled by increases in pERK5, PSD95, and pSyn levels (Fig. 3d-f). The compound promoted only a modest increase in pTrkB levels in 3xTg-AD^HFD^ mice (Fig. 3c). Two considerations can explain this result: 1) transgenic AD mice showed impaired TrkB signaling despite the presence of elevated BDNF levels ^62^; 2) drug-related trophic effects can, at least in part, be mediated by BDNF-independent mechanisms ^26,63^. In line with our previous results ^26^, exenatide was found to be effective in reducing proBDNF/p75NTR signaling, a mechanism that occludes the activation of the pro-apoptotic pJNK and pERK_1,2_ molecules (Fig. 4).

In 3xTg-AD^HFD^ mice, exenatide was found to modulate neuroinflammation as shown by the drug-driven reduction of NF-κB levels (Fig. 5a). This finding is supported by several studies showing an anti-inflammatory role for GLP-1R agonists in neurodegenerative conditions ^28,63^. The drug-mediated reduction of NF-kB was mirrored by a net decrease in PPARα and PPARγ levels (Fig. 5b-d). This result supports the idea that exenatide, by blocking inflammatory signaling, prevents the activation of the endogenous anti-inflammatory responses ^43^.

In line with our previous findings ^25^, exenatide had no effect on Aβ and p-tau pathology (Fig. 2a-d).

### Effect of exenatide on cognition

Our findings converge towards potential pro-cognitive effects of exenatide. However, the analysis of the behavioral data failed to reveal significant changes in the memory performances of 3xTg-AD mice. These results are not in line with reports indicating detrimental cognitive effects of HFD in mice ^34,64^. A series of compensatory mechanisms may have produced the lack of cognitive impairment in the study animals. One issue concerns the fact that our 12-month-old (m.o.) 3xTg-AD mice fail to display cognitive deficits. MWM and NOR performances of vehicle-treated 3xTg-AD^CD^ mice did overlap with those observed in age-matched wild-type animals employed by our lab in previous studies [cfr. ^26,31^, and data not shown], thereby supporting the notion that our 12-m.o. 3xTg-AD mice were still in a pre-symptomatic phase. Interestingly, this lack of cognitive deficits occurred even in the presence of overt signs of Aβ and p-tau pathology (Fig. 2). This result is not surprising as previous findings showed that cognitive performances of the 3xTg-AD mice could be restored or preserved even without decreasing Aβ or p-tau levels, thereby indicating the presence of amyloid-independent compensatory mechanisms ^30,65,66^. In addition, these mice were able to cope with the HFD-driven metabolic insult. 3xTg-AD^HFD^ mice showed increased peripheral levels of insulin (Fig. 1i) as indicated by the absence of signs of hyperglycemia or diabetes in the study group (Fig. 1c-d and 1h). As insulin actively modulates cognition, it is conceivable that, in the CNS of the 3xTg-AD^HFD^ mice, the enhanced hormone levels may have worked towards the facilitation of a neurotrophic drive that counteracted the impaired BDNF-TrkB axis (Fig. 3a and 3c). This possibility is indirectly indicated by the decreased phosphorylation, and therefore the increased activation, of IRS-1 that we have found in the brain of the 3xTg-AD^HFD^ animals (Fig. 3g), a phenomenon that supports the idea of an ongoing neurotrophic activity exerted by insulin. Lending further support to this hypothesis, the exenatide-driven recovery of the BDNF/TrkB cascade was paralleled by an increase in pIRS-1 levels (Fig. 3), thereby indicating the potential occlusion of any additional insulin-mediated neurotrophic effect.

Exenatide also failed to produce behavioral changes in 3xTg-AD^CD^. The finding was unexpected as the molecule has been described to exert a pro-cognitive activity in WT animals ^26^. A possible explanation for this result comes from recent evidence showing that, to promote cognitive improvement in an AD mouse model, BDNF-signaling requires adult neurogenesis ^67^. Twelve m.o. 3xTg-AD animals have been shown to exhibit signs of impaired hippocampal neurogenesis ^68,69^, thereby preventing the beneficial role of exenatide-driven BDNF signaling.

## Conclusions

Despite significant efforts in the field of AD-related therapy, no disease-modifying drugs are currently available. A reconsideration of the “amyloid hypothesis” as well as a more complex view of the disease and the synergistic role played by additional mechanisms is animating the AD community ^4,64,70^. Growing evidence indicate that the modulation of neurotrophic signaling may be a promising therapeutic avenue to be explored in neurodegenerative conditions and AD ^71,72^. Our findings support the need to investigate further exenatide in clinical trials targeting MCI subjects or patients suffering from the early stage of dementia.

## Supporting information

## Acknowledgments

The authors thank all the members of the Molecular Neurology Unit for helpful discussions. The authors are in debt with Paola Siccu and Annalisa Nespoli for technical assistance with IHC.

## Author contribution

SLS conceived, designed, and supervised the study. MB performed *in vivo* treatment, metabolic assays, behavioral testing, and sample collection. VC and AC performed and analyzed WB data. RL performed and analyzed IHC data. MB and AG analyzed and interpreted the data. MB, AG, and SLS wrote the manuscript. All authors approved the final version of the manuscript.

