## Supplementary material for "Exenatide reverts the high-fat-diet-induced impairment of BDNF signaling and inflammatory response in an animal model of Alzheimer’s disease"

**Supplementary Figure 1**

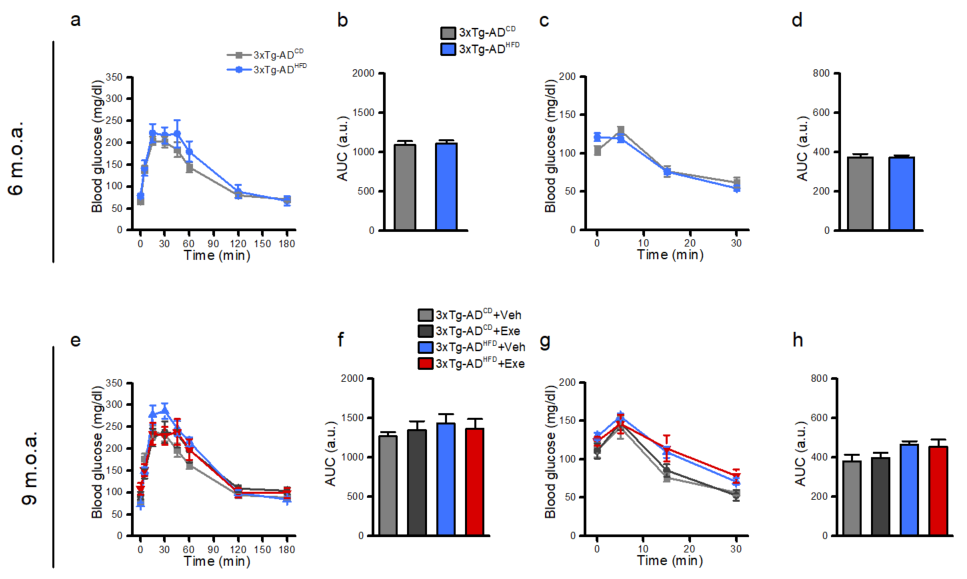

**Supplementary figure 1. Metabolic changes of 3xTg-AD mice at 6 and 9 m.o.a.** (a) Graphs depict the intra-peritoneal GTT (glucose tolerance test) curve of 6 m.o. mice performed at the beginning of the HFD or control diet treatments. (b) Bar graphs illustrate the GTT quantifications expressed as area under the curve (AUC). (c) Graphs illustrate the intra-peritoneal ITT (insulin tolerance test) of the 3xTg-AD^CD^ and 3xTg-AD^HFD^ mice. (d) Bar graphs illustrate the ITT quantifications expressed as AUC. (e) Graphs depict the intra-peritoneal GTT curve of 9 m.o. mice at the beginning of the exenatide or control treatments. (f) Bar graphs illustrate the GTT quantifications expressed as AUC. (g) Graphs illustrate the intra-peritoneal ITT of the four study groups. (h) Bar graphs illustrate the ITT quantifications expressed as AUC. Means were compared by one-way ANOVA followed by Tukey post-hoc test. No statistically significant differences were observed among the study groups at 6 and 9 m.o.a. Please, note that 9 m.o. 3xTg-AD^HFD^ mice start to develop early signs of insulin resistance. The phenomenon, however, does not reach statistical significance. Abbreviations: Veh, vehicle; Exe, exenatide.

**Supplementary Table 1**

| Antibody | Dilution | Supplier | Catalog # |
| --- | --- | --- | --- |
| Anti-pSynapsin | 1:1000 | Cell signaling | 2311S |
| Anti-Synapsin | 1:1000 | Invitrogen | A6442 |
| Anti-BDNF | 1:1000 | Abcam | Ab108319 |
| Anti-proBDNF | 1:500 | Millipore | MABN110 |
| Anti-TrkB | 1:1000 | Abcam | Ab18987 |
| Anti-pTrkB | 1:500 | Abcam | Ab109684 |
| Anti-p75NTR | 1:20000 | Abcam | Ab52987 |
| Anti-pERK1/2 | 1:500 | Santa Cruz | Sc-7383 |
| Anti-ERK1/2 | 1:200 | Santa Cruz | SC-94 |
| Anti-pERK5 | 1:1000 | Cell signaling | 3371S |
| Anti-ERK5 | 1:1000 | Cell signaling | 3372s |
| Anti-pJNK | 1:500 | Santa Cruz | Sc-12882-R |
| Anti-JNK | 1:200 | Santa Cruz | Sc-571 |
| Anti-PSD95 | 1:1000 | Abcam | Ab18258 |
| β-actin | 1:10000 | Cell Signalling | 12262S |
| Anti-NF-κB | 1:500 | Santa Cruz | Sc-372 |
| Anti-PPARγ | 1:500 | Abcam | 209350 |
| Anti-PPARβ/δ | 1:500 | Invitrogen | PA529678 |
| Anti-PPARα | 1:500 | Abcam | 8934 |
| Anti-IRS1 | 1:1000 | Cell signaling | 3407S |
| Anti-pIRS1 | 1:1000 | Cell signaling | 2385s |
| Anti-pCREB | 1:1000 | Cell signaling | 9198s |
| Anti-CREB | 1:1000 | Cell signaling | 4820s |
